# Transcriptomic evaluation of immune-infiltrated patient-derived tumor organoids as preclinical models in renal cell carcinoma

**DOI:** 10.64898/2026.01.28.699468

**Authors:** Liangwei Yin, Léonard Lugand, Jules Russick, Joel Lemaoult, Christophe Battail

**Affiliations:** Université Grenoble Alpes, CEA, INSERM, UA13 BGE, CEA, Grenoble, France; Hemato-Immunology Research Department, CEA, DRF-Francois Jacob Institute, Saint-Louis Hospital, Paris, France - INSERM UMR1342, IRSL, Paris University, Paris, France

**Keywords:** clear cell renal cell carcinoma, patient-derived tumor organoids, tumor infiltrating lymphocytes, transcriptomics, immunotherapy response

## Abstract

Patient-derived tumor organoids (PDTOs) have emerged as valuable preclinical models for studying tumor biology and therapeutic responses. However, despite the development of protocols for tumor dissociation and immune cell infiltration, their fidelity in representing clear cell renal cell carcinoma (ccRCC) tumors remains poorly characterized. To address this, we established matched samples, including tumor tissue, enzymatically dissociated tumor, formed PDTOs, and immune-infiltrated PDTOs from three ccRCC patients, and performed bulk RNA sequencing to capture dynamic molecular changes across the experimental workflow. Our analyses revealed that tumor dissociation triggered stress-related transcriptional changes, marked by the upregulation of heat shock genes (e.g., HSPA1A) and the downregulation of the hypoxia pathway, while PDTOs recapitulated hypoxic signaling. Immune-infiltrated PDTOs retained critical immune signatures including T-effector, and exhibited an enhanced pro-inflammatory phenotype (CXCL10, JAK-STAT). Furthermore, predictive gene signatures and immunotherapy response scores further underscored the clinical relevance of immune-infiltrated PDTOs consistent with the original tumor tissue. Collectively, these findings validate immune-infiltrated PDTOs as robust, patient-specific models for personalized therapeutic exploration, offering a platform to optimize immunotherapy strategies in ccRCC.

## Introduction

Clear cell renal cell carcinoma (ccRCC) is the most common subtype of renal cancer, characterized by a highly heterogeneous tumor microenvironment that drives tumor progression and confers resistance to therapies^1,2^. While surgical treatment remains the standard for localized ccRCC, metastatic ccRCC relies on systemic therapies, including combinations of tyrosine kinase inhibitors (TKIs) and immune checkpoint blockade (ICB) or dual ICBs, as strongly recommended by the EAU guidelines^3^. Despite the advancements in immunotherapy, treatment outcomes remain suboptimal in terms of variable patient response rates between 7-59%^4–6^. This variability underscores therapeutic challenges posed by tumor heterogeneity and complex resistance mechanisms, which have hindered the identification of reliable molecular biomarkers for predicting therapeutic efficacy^7^. To address these limitations, patient derived tumor organoids (PDTOs) have emerged as promising clinical tools for precision oncology, in drug screening, treatment optimization, and outcome predictions^8–10^.

Conventional preclinical models, such as two-dimensional culture of cell lines or tumor cells, inadequately replicate the physical structure and tumor microenvironment (TME)^11^. These systems often fail to preserve hypoxia gradients, immune cell activity, and tumor-immune cells interactions, which are essential for modeling immunotherapy responses. Recently, a standardized protocol of patient derived tumor organoids (PDTOs) with infiltrated lymphocytes has been recently established to retain the original tumor complexity, in terms of tumor cell heterogeneity and TME characteristics^12^. These immune-infiltrated PDTOs (infiltrated-PDTOs) offer a transformative platform to study immunotherapy mechanisms and resistance in a clinically relevant setting.

However, it remains to be further characterized for the artifacts of experimental steps in the formation of organoids, and the representativeness of formed organoids in the original tumor. For examples, tumor dissociation disrupted the physical structure of the original tumor^13^, which may impact the composition of formed organoids. The re-infiltration of immune cells, also raised of the question on how the organoids mimicked TME and modeled immune evasion of the original tumor. Critically, transcriptomics profiles of experimental samples may indicate the extent of the resemblance between the original tumor and matched organoids, and immune checkpoint genes could exhibit the fidelity of organoids in preclinical trials of immunotherapy.

Therefore, we evaluate the fidelity and translational potential of infiltrated-PDTOs as preclinical models for immunotherapy in ccRCC. Using a standardized protocol, we generated matched patient samples including primary tumor tissue, dissociated tumor cells, PDTOs, and infiltrated-PDTOs from three ccRCC patients. Through transcriptomic profiling, we characterized molecular and cellular changes during tumor dissociation and organoids formation, identifying differentially expressed genes, transcription factors, and pathways. We further validated infiltrated-PDTOs using tumor- and immune-specific signatures, and assessed their suitability as preclinical models by correlating organoid responses with clinical predictive biomarkers. By bridging the gap between in vitro models and clinical complexity, this work establishes infiltrated-PDTOs as a robust platform for advancing personalized immunotherapy in ccRCC.

## Materials and Methods

### Tumor collection

Fresh tumor specimens were obtained from patients undergoing nephrectomy for localized clear cell renal cell carcinoma (ccRCC) (Saint-Louis Hospital, Paris, France) (Table 1). These specimens, classified as surgical waste, were collected with the patients’ written informed consent prior to inclusion in the study. Tumors were collected by a pathologist under sterile conditions, and immediately transferred to Roswell Parks Memorial Institute (RPMI, Gibco) medium, supplemented with antibiotics on ice for transport to the laboratory within 30 min of excision.

**Table 1.** Clinical information of patients with kidney cancer.

| Patient ID | ISUP grade | PD-L1 CPS | Age | Sex | TNM |
| --- | --- | --- | --- | --- | --- |
| Patient 1 | 2 | Positive | 65-70 | F | pT3aNxR0 |
| Patient 2 | 4 | Positive | 75-80 | F | pT3aNxR0 |
| Patient 3 | 2 | Positive | 65-70 | M | PT1aNxR0 |

### Tumor digestion

Tumor samples were finely minced into ∼1 mm fragments and suspended in 2 mL Roswell Park Memorial Institute (RPMI) medium (Gibco, Thermo Fisher Scientific) at the bottom of a Petri dish. Three replicates of 100 ul each were aliquoted into a U-bottom 96-well plate (labeled “tumor tissue”) and stored on ice until RNA extraction.

The remaining minced tumor material was digested in 6 mL Hank’s Balanced Salt Solution with calcium and magnesium (HBSS; Gibco, Thermo Fisher Scientific), containing 0.125 WU/mL LiberaseTM (Sigma-Aldrich) and 100 U/mL DNase I (Sigma-Aldrich), for 35 min at 37 °C in a water bath. The digested suspension passed through a 100 µm cell strainer to remove undigested debris, and the reaction was stopped by adding 40mL cold RPMI, supplement with 10% fetal bovine serum (FBS; Sigma-Aldrich). Cells were pelleted by centrifugation at 400 ×g for 5 min. The red blood cells were lysed using RBC Lysis Buffer (BioLegend) according to the manufacturer’s instructions.

Three 100 µL aliquots of the dissociated tumor cell suspension were transferred into the same 96-well plate (labeled “dissociated tumor”). For separation of tumor and immune cells, magnetic-activated cell sorting (MACS) was performed using CD45 MicroBeads (Miltenyi Biotec) according to the manufacturer’s instructions.

### RNA extraction for Bulk RNA Barcoding and sequencing (BRB-seq)

RNA extraction from both “tumor tissue” and “dissociated tumor” samples was performed using the Organoid DRUG-seq Kit (Alithea Genomics) according to the manufacturer’s instructions. Extracted RNA was stored in the provided 96-well plate at −80 °C until shipment.

### Patient-Derived Tumor Organoid (PDTO) formation

Non-immune (CD45^−^) and immune (CD45^+^) fractions were collected after MACS separation. CD45^−^ cells were rinsed and resuspended in Renal Cell Carcinoma Medium (RCCM), consisting of Dulbecco’s Modified Eagle Medium (DMEM; Gibco) and Ham’s F-12 medium (Gibco) mixed at a 3:1 ratio (v/v), supplemented with 10% FBS, 2 mM L-glutamine (Gibco), 100 U/mL penicillin–streptomycin (Gibco), 10 ng/mL epidermal growth factor (EGF; Sigma-Aldrich) and 0.4 µg/mL hydrocortisone (Sigma-Aldrich).

For organoid formation, 10 000 CD45^−^ cells were seeded in 100 µL RCCM into a 96-well cell-repellent plate (Greiner Bio-One) and centrifuged at 500 ×g for 1 min to promote aggregation. In parallel, CD45^+^ cells were activated in a 96-well U-bottom plate (30 000 cells/well) with 40 ng/mL IL-15 (PeproTech).

After 48 h, activated CD45^+^ cells from 10 wells were transferred to 10 wells containing formed PDTOs, achieving a 3:1 ratio (30 000 immune cells per 10 000 CD45^−^ cells initially seeded). Another 10 PDTO wells were left untreated.

After 24 h, all PDTOs (labeled “PDTOs”) and co-cultured PDTOs with immune cells (labeled “infiltrated PDTOs”) were washed three times in PBS to remove non-infiltrated immune cells. RNA extraction was then performed in each well using the Organoid DRUG-seq Kit (Alithea Genomics) as described above. Extracted RNA was added to the Alithea sample plate and stored at −80 °C until shipment.

### Library construction and sequencing

Extracted RNA from samples described above was sent to Alithea Genomics SA (Lausanne, Switzerland) for library preparation and sequencing using extraction-free, highly multiplexed, 3′-end bulk RNA barcoding and sequencing (MERCURIUSTM Organoid DRUG-seq service). Bulk RNA Barcoding and sequencing (BRB-seq) libraries were prepared using the MERCURIUSTM Organoid DRUG-seq library preparation kit for Illumina, according to the manufacturer’s manual (Alithea Genomics, #10861). The constructed libraries were sequenced on an Illumina NovaSeq 6000 platform.

RNA-seq reads generated from BRB-seq were then aligned and quantified using STAR (version 2.7.9) against the GRCh38 HUMAN reference genome and quality control was performed using FastQC (version 0.11.9) and RSeQC (version 4.0.0)^14–16^. Unique molecular identifiers (UMIs) counts per gene were used for downstream analyses. For sample with biological duplicates, duplicates were excluded if they had a lower number of detected genes (CPM: counts per million > 0) or a Pearson correlation coefficient below 0.4 compared to other duplicates.

### Transcriptional analysis

Differential gene expression (DGE) analysis, Transcription factors (TF) analysis, and pathway activity (PA) analysis were performed and visualized using decoupleR (python version 1.8.0)^17^. Briefly, raw reads count of samples were used as the input and genes were excluded if number of their reads was less than 10. Differential expressed genes (DEGs) were inferred with the embedded Deseq2, analyzing the statistics of genes between two conditions^18^. DEGs were taken as significant genes, if fold changes of DEGs were greater than 2 with an adjusted p-value less than 0.05. TF activity inference was conducted using the univariate linear model, with the statistics results of DEGs and a CollecTRI network database (The regulations between TF and genes were included)^19^. Pathway activity was inferred using multivariate linear model on the statistics results of DEGs and a PROGENy resource (14 curated pathways, containing weights for the interaction between genes and pathways)^20^.

In addition, over representation analysis (ORA) and gene set enrichment analysis (GSEA) were also performed using gene ontology (GO) terms and curated KEGG (Kyoto Encyclopedia of Genes and Genomes) pathways from Molecular Signature Database (MsigDB)^21^.

### Transcriptional signature estimation

For signature analysis, raw read counts were normalized to transcripts per million (TPM) and log_2_-transformed (*log*_*2*_ *(TPM + 1)*). Molecular signature analysis was conducted using ccRCC-specific subset signatures collected from the paper of Motzer et al (Signatures: T effector, angiogenesis, myeloid Inflammation signatures, cell cycle, stroma, the complement cascade, small nucleolar RNAs (snoRNAs), and metabolism-related pathways, including fatty acid oxidation (FAO)/AMP-activated protein kinase (AMPK) signaling, fatty acid synthesis (FAS)/ pentose phosphate, and biological oxidation pathways)^22^.

Tumor signatures were also assessed by the ESTIMATE R package (Estimate of Stromal and Immune Cells in Malignant Tumor Tissues from Expression Data), including stromal, immune, tumor purity and ESTIMATE scores^23^.

### Tumor immune infiltration analysis

Tumor immune infiltration levels were evaluated by single sample gene set enrichment analysis (ssGSEA), with immune-related gene sets derived from a pan-cancer study^24^. Calculations were performed with GSVA using the ssGSEA algorithm^25^.

### Immunotherapy response prediction

To predict patients’ response to immunotherapy, TIDE (Tumor Immune Dysfunction and Exclusion) scores were computed based on the gene expression profiles of our samples using the website online: http://tide.dfci.harvard.edu/^26^. Based on TIDE output, samples were classified as predicted responders or non-responders. In parallel, expression levels of canonical immune checkpoint genes were directly compared across samples.

## Results

### Generation of infiltrated-PDTOs from ccRCC patients

To derive patient-derived tumor organoids (PDTO), a tumor organoids model has been established and evaluated in the context of immunotherapies^12^. We obtained tumor samples from three patients (Patient 1, Patient 2, and Patient 3) (Table 1), and generated organoids following the established protocol^12^, including tumor dissociation, selection of non-immune fraction and immune fraction, PDTO formation, and PDTO with infiltrating lymphocytes (infiltrated-PDTO) (Figure 1).

**Figure 1.**
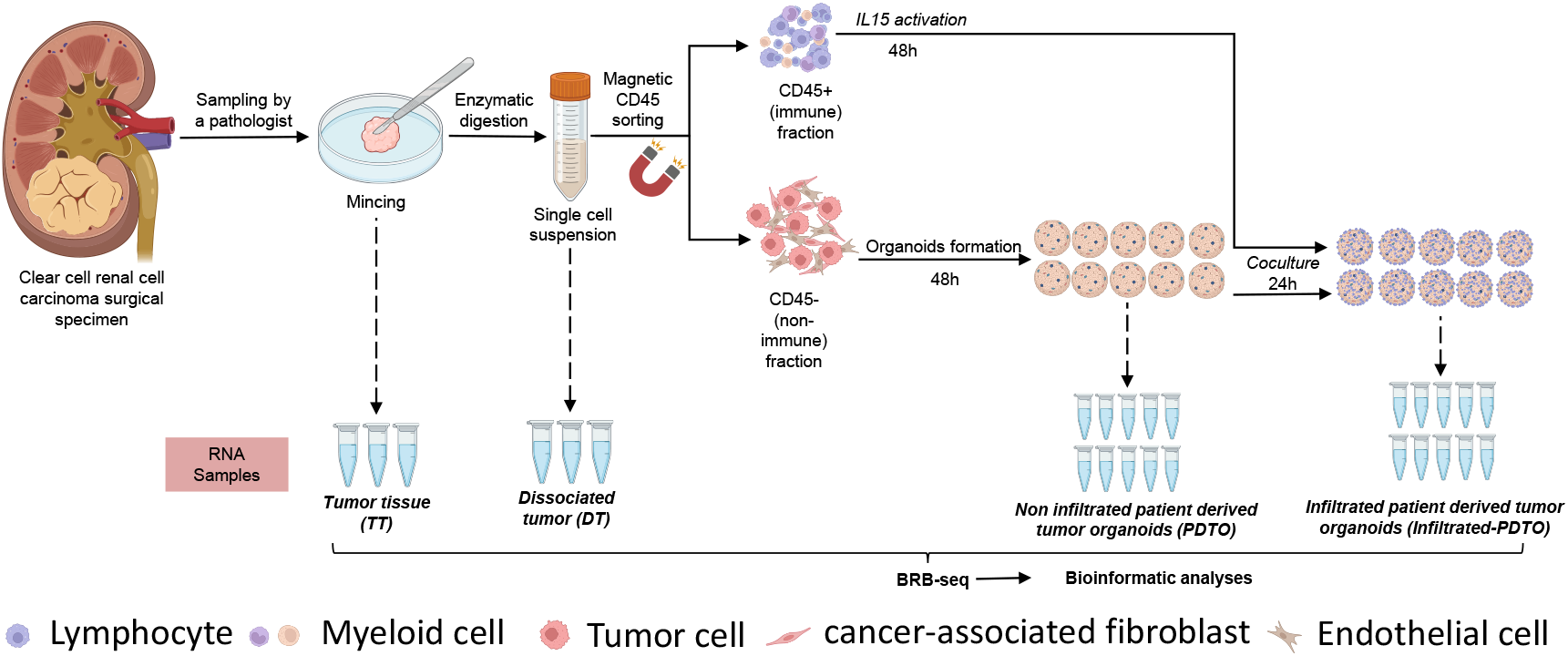
Experimental design for the establishment of patient-derived tumor organoids (PDTO), and patient-derived tumor organoids with infiltrating lymphocytes (infiltrated-PDTO).

Tumor tissue was firstly dissociated into fragments, homogenized, and enzymatically digested (dissociated tumor). Then, single-cell suspensions were separated into CD45^−^ (tumor-enriched) and CD45^+^ (immune cells) fractions using magnetic-activated cell sorting (MACS) with anti-CD45 microbeads. For PDTO generation, CD45^−^ cells (10,000/well) were seeded in cell-repellent 96-well plates with Renal Cell Carcinoma Medium (RCCM), centrifuged (500 × g, 1 min), and cultured for 48 hours to form organoids. Concurrently, CD45^+^ cells were activated with 30 ng/mL IL-15 in complete RPMI for 48 hours. Activated tumor-infiltrating lymphocytes (TILs, 30, 000 cells) were co-cultured with spheroids (a ratio of tumor to immune cells: 1:3) for 24 hours, while control spheroids remained untreated. Post-co-culture, organoids were washed, lysed, and processed for RNA extraction to profile infiltrated and non-infiltrated models.

Based on sequencing data, most of samples were kept for transcriptional analysis, except 1 PDTO and 1 infiltrated-PDTO in Patient 1, and 1 PDTO in Patient 2 (Table S1). The filtration was based on the Pearson correlation between samples inside their category, the number of expressed genes, and principal component analysis (Figure S1-2). Using the transcriptional profile, we then evaluated the fidelity of infiltrated-PDTOs as preclinical models in ccRCC for treatment response.

### Molecular characterization of tumor dissociation and PDTO formation

To explore how tumor dissociation and cell culture influence PDTO formation, we set out to investigate differences in gene expression, transcription factors (TFs), and cancer pathways. Differential gene expression (DGE) analyses were performed for two comparisons: tumor tissue versus dissociated tumor (effect of enzymatic dissociation), and dissociated tumor versus PDTO (effect of PDTO formation). In our representative case (Patient 1), 18,131 genes were kept based on their total reads count across all samples. During tumor dissociation, 1,980 genes were significantly up-regulated and 1,482 genes were down-regulated in dissociated tumor relative to tumor tissue (fold change > 2; adjusted p-value < 0.05; Figure 2A; Table S2). Out of top differentially expressed genes (DEGs), *RGS2, HSPA8, HSPA6*, and *ATF3* were canonical stress-responsive genes, triggered by dissociation stress^27,28^. Consistently, inference of TFs activity revealed activated TFs such as HSF1 (Heat shock transcription factor 1), RELA, NFKB1, and JUN in the dissociated tumor (Figure 2A), mediating cellular responses to enzymatic dissociation. Moreover, analysis of canonical cancer pathways indicated an active EGFR pathway alongside an inactive hypoxia pathway, indicating cell proliferation and reverse low oxygen of dissociated tumors (Figure 2A). No significant pathway was found using gene set enrichment analysis (GSEA) (Table S3). In the progression from dissociated tumor to PDTO, 1,086 up-regulated and 1,920 down-regulated DEGs were identified in PDTO (fold change > 2; p-value < 0.005; Figure 2B). The p-value was stricter due to imbalance of samples (Figure S3). Top DEGs such as *HLA-DRB1*, and *HLA-DPA1* encode components of the MHC class II complex, likely expressed by immune cells, highlighting the depletion of immune cells in PDTO. Inactive TFs such as RFXAP, RFXANK, and RFX5 were inferred (Figure 2B), which were essential for activating MHC class II genes. Relative to dissociated tumor, hypoxia was restored, and activation of the PI3K and MAPK pathway indicated the proliferation of tumor cells in PDTO (Figure 2B). The suppression of the JAK-STAT pathway further highlights the depletion of immune components in PDTO, which was also validated by the down-regulation of KEGG pathways such as T cell receptor signaling pathway and chemokine signaling pathway (Table S3).

**Figure 2.**
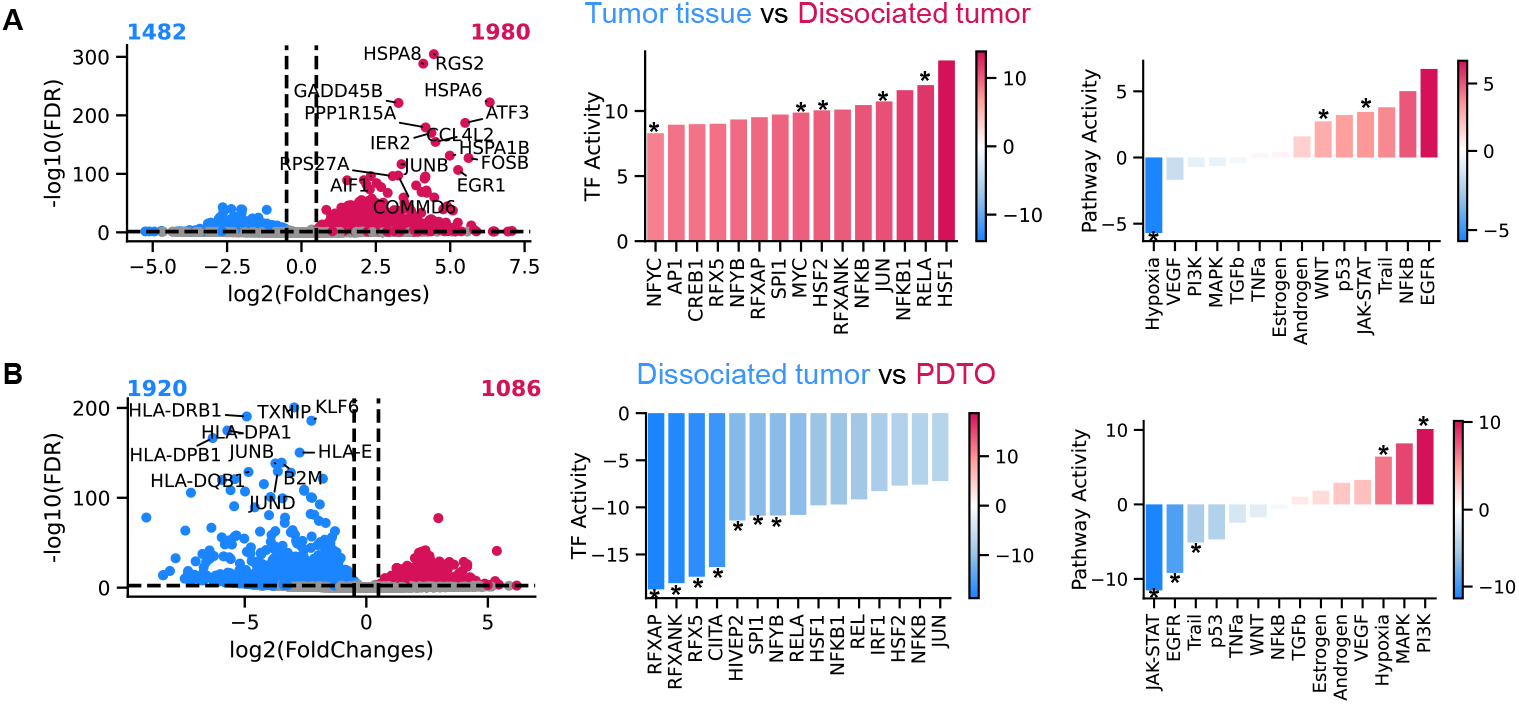
Differential gene expression, transcription factor, and pathway activity analysis in tumor dissociation, and the formation of PDTO for Patient 1. A. Comparisons between tumor tissues to dissociated tumor. B. Comparisons between dissociated tumors and PDTOs. Red color indicated the upregulation, and blue color indicated the downregulation. Left plots were volcano plots of differential expressed genes (DEGs), middle plots were of TF activity inferred from DEGs, and right plots were of pathway activity inferred from DEGs. Noted that “*” refers to shared TFs and pathways across three patients.

In Patient 2 and Patient 3, fewer DEGs were detected in tumor dissociation and PDTO formation, relatively compared to Patient 1 (Figure S4-5; Table S4-5). In tumor dissociation, the upregulation of ribosome related genes and heat shock genes (*RPL37A, HSPA8*) was highlighted for both patients. Consistent with stress response and cell proliferation genes, Patient 2 exhibited inactive TFs (SP53, ARNT) and activation of the JAK-STAT pathway, and Patient 3 was characterized with an active PI3K pathway (Figure S4A-5A). The downregulation of hypoxia pathway was present in all patients, introducing a bias of the hypoxic microenvironment for dissociated tumor^29^. In PDTO formation, despite different DEGs, inactive TFs (RFXAP, RFXANK), active PI3K, inactive JAK-STAT pathway, and down-regulation of immune related KEGG pathways were observed in PDTO from all three patients (Figure S4B-5B; Table S6-7). Notably, the hypoxic microenvironment was restored in the formation PDTO from tumor dissociation (Figure 2B, Figure S4B-5B).

To investigate the technical effects of experimental procedures, we identified common DEGs across three patients for these two comparisons, thereby minimizing patient-specific tumor heterogeneity. In tumor dissociation, we identified 90 upregulated and 165 downregulated DEGs common to all patients (Figure S6A; Table S2,4-5). Gene ontology (GO) analysis reported that up-regulated genes were enriched in processes such as protein refolding, cytosolic ribosome, and C3HC4 type ring finger domain binding (Figure S6A), suggesting stress response activation. Down-regulated genes were associated with the ERBB3 signaling pathway, and xenobiotic transmembrane transporter (Figure S6A), suggesting the loss of tumor 3D structure. In PDTO formation, 58 up-regulated DEGs were enriched in GO terms such as negative regulation of granulocyte chemotaxis and cyclooxygenase pathway, and 233 down-regulated DEGs were mainly associated with the MHC protein related process, emphasizing the depletion of immune cells in PDTO (Figure S6B).

Collectively, our analysis highlights the heterogeneity and commonality at transcriptional level during tumor dissociation, and PTDO formation for patient derived tumor. While individual patients exhibited varying number of DEGs and dysregulated TFs and pathways, several pathways as hypoxia, PI3K and JAK-STAT emerged, underscoring oxidative stress and cell proliferation in tumor culture and PDTO development.

### Molecular differences between tumor tissue, PDTO, and infiltrated-PDTO

In order to investigate the molecular distinction of infiltrated-PDTO, we next performed DGE analysis directly between tumor tissue and PDTO or infiltrated-PDTO. In Patient 1, PDTOs exhibited 1,799 up-regulated and 2,204 down-regulated DEGs compared to tumor tissue, while infiltrated-PDTO showed 2,579 up-regulated and 2844 down-regulated genes relative to tumor tissue (fold change > 2, p-value < 0.005; Table S2). Integrated with DEGs between dissociated tumor and tumor tissue, a heatmap of 7747 DEGs illustrates these differences (Figure 3A).

**Figure 3.**
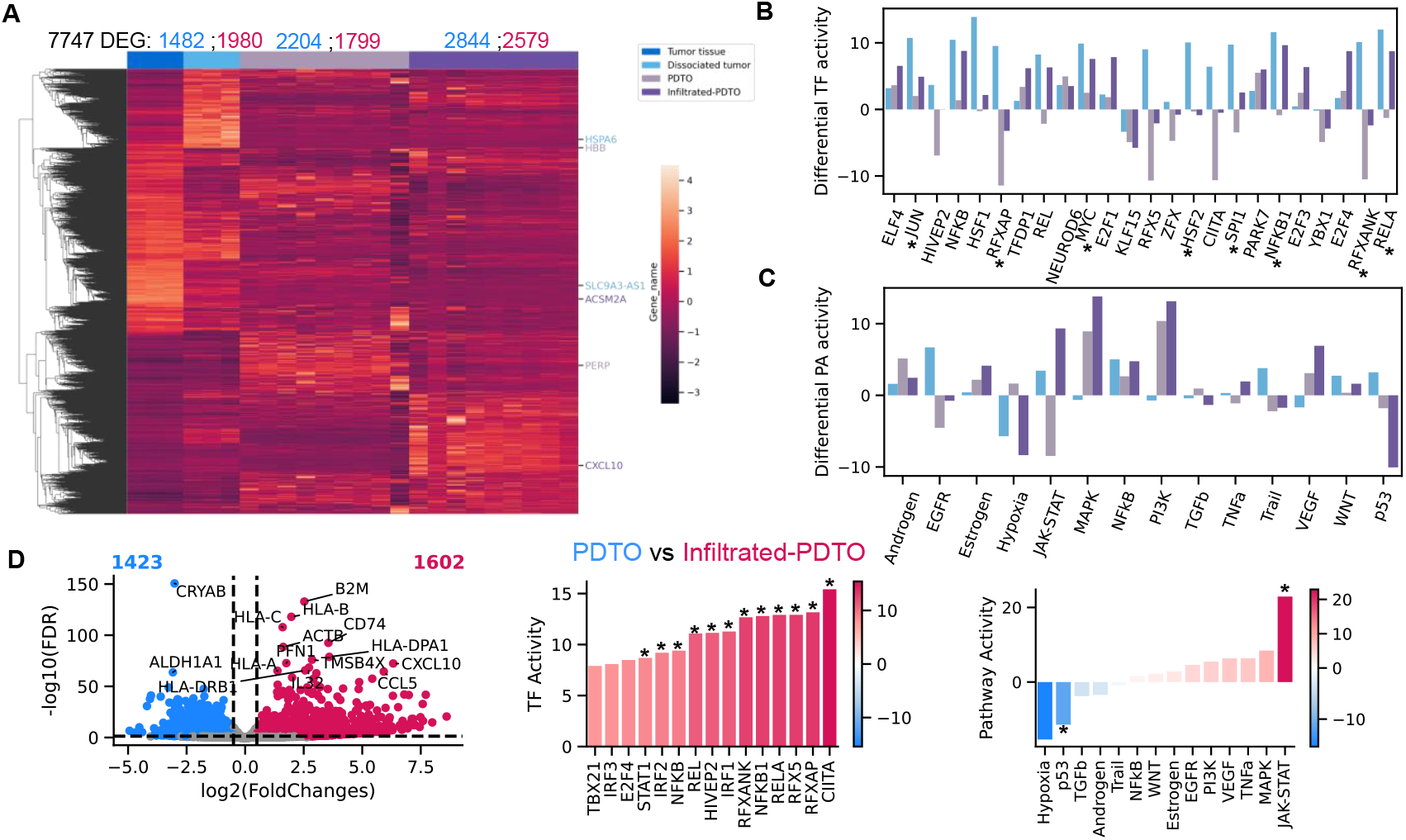
Differential gene expression, transcription factor, and pathway activity analysis in the formation of PDTO for Patient 1. A. A heatmap for all DEGs from the comparison between tumor tissue and dissociated tumor or PDTO or infiltrated-PDTO. B. TF analysis for the comparison between tumor tissue and dissociated tumor or PDTO or infiltrated-PDTO. Top 10 significant TFs were selected for three comparisons, and 24 TFs were obtained. C. PA analysis for the comparison between tumor tissue and dissociated tumor or PDTO or infiltrated-PDTO. D. Comparisons between PDTO vs infiltrated-PDTO. Noted that “*” refers to shared TFs and pathways across three patients.

Compared to tumor tissue, several TFs (ELF4, JUNE, etc.) were consistently dysregulated in dissociated tumor, PDTO, and infiltrated-PDTO (Figure 3B). While PDTO demonstrated inactive TFs such as ZFX, RFXANK, RFX5, CIITA, and RFXAP, likely due to the absence of immune cells, infiltrated-PDTOs showed activation of E2F3, ELF4, TFDP1, and E2F1 (cell cycle TFs), suggesting tumor cells remained proliferative after the re-infiltration of immune cells. Furthermore, pathway analysis indicated the absence of immune cells by active hypoxia and TGFb pathway, alongside the inactive JAT-STAT and TNFa pathways in PDTO. In contrast, infiltrated-PDTOs exhibited activation of JAK-STAT, PI3K, VEGF pathway, with an inactive p53 pathway, indicating cytokine signaling and interactions between tumor cells and immune cells. GSEA also revealed the enrichment of downregulation of immune cells related KEGG pathway in PDTO, and downregulation of metabolism related pathways in infiltrated-PDTO (Table S3)

For Patient 2 and Patient 3, PDTOs had 1,529 up-regulated, and 3,357 down-regulated DEGs, and 340 up-regulated, 1,269 down-regulated DEGs, respectively (Figure S7-8A; Table S4-5). Infiltrated-PDTO showed 1,357 up-regulated, and 3,068 down-regulated genes, and 274 up-regulated, 1,716 down-regulated genes, respectively (Figure S7-8A). All DEGs were illustrated in the heatmap (Figure S7-8B). Both patients displayed inactive MHC class II related TFs and inactive JAK-STAT pathway in PDTO, and a highly active JAK-STAT pathway in infiltrated-PDTO (Figure S7-8CD).

Common DEGs across all patients included 190 up-regulated and 937 down-regulated genes in PDTO, and 138 up-regulated genes and 726 down-regulated genes in infiltrated-PDTO (Figure S9). GO analysis highlighted enrichment of MHC-related and immune cell-related terms among down-regulated genes in PDTO, and enrichment of small nuclear ribonucleoproteins (snRNPs) among up-regulated genes in infiltrated-PDTO. Upregulation of snRNPs were associated with immune evasion^30^. This was complemented by down-regulated KEGG pathways such as hematopoietic cell lineage and T cell receptor signaling pathway in PDTO and down-regulation of metabolism and signaling pathways (e.g., Alanine aspartate and glutamate metabolism, MAPK signaling pathway) in infiltrated-PDTO.

The modest number of DEGs (90 upregulated and 165 downregulated) between tumor tissue and dissociated tumor suggests minimal transcriptomic alterations due to dissociation. In contrast, PDTOs exhibited a substantial number of DEGs (190 upregulated and 937 downregulated), while infiltrated-PDTOs had fewer DEGs (138 upregulated and 726 downregulated) compared to tumor tissue. This pattern indicates that the presence of infiltrating immune cells enhances the transcriptomic resemblance of PDTOs to the original tumor tissue.

### Comparisons between PDTO and infiltrated-PDTO

To further investigate the impact of immune fraction on PDTO, we conducted a comparative transcriptomic analysis between PDTO and infiltrated-PDTO. In Patient 1, we identified 1,602 up-regulated and 1,423 down-regulated genes in infiltrated-PDTO relative to PDTOs (Figure 3D, left panel; p value < 0.05, fold change > 2; Table S3). TFs and pathway analyses revealed highly active TFs such as CIITA, RFXAP, etc. and activation of the JAK-STAT pathway (Figure 3D, right panel), suggesting tumor-immune interaction in infiltrated-PDTO.

Although Patients 2 and 3 exhibited different numbers of DEGs (396 up-regulated and 148 down-regulated for Patient 2; 337 up-regulated and 337 down-regulated for Patient 3; Figure S10; Table S4-5), notably, there was increased activity of CIITA, IRF1, and RELA, along with heightened activity of the JAK-STAT pathway (Figure S10). Based common genes and pathways across patients, GO analysis reported the enrichment of MHC related biological function (Figure S11), and GSEA indicated the up-regulation of cytokine-cytokine receptor interaction, antigen processing and presentation, and natural killer cell mediated cytotoxicity.

While only a small subset of DEGs was common across all three patients (155 up-regulated and 36 down-regulated), the consistent enrichment of immune-related pathways underscores the specificity of immune cell infiltration effects in infiltrated-PDTOs compared to PDTOs.

### Tumor and immune signatures of infiltrated-PDTO

The re-infiltration of immune fraction into PDTO plays an important role in recapitulating tumor immune microenvironment of the original tumor. Hence, we analyzed specific tumor and immune signatures across our categories of samples. Building upon previously established pathway signatures of molecular subsets in ccRCC (T effector, Cell Cycle, FAS(fatty acid synthesis)/pentose phosphate, Complement Cascade, Omega Oxidation, Myeloid Inflammation, Angiogenesis, FAO/AMPK(fatty acid oxidation/ AMP-activated protein kinase), Stroma, snoRNA)^22^, we characterized our samples from tumor tissue to infiltrated-PDTO (Figure 4A). In Patient 1, tumor tissue was characterized with the high level of angiogenesis and complement cascade. Dissociated tumor exhibited elevated T effector and myeloid inflammation, with the decrease in other signatures. PDTO showed a significant decline in T effector, while other signatures remained similar to dissociated tumor. Notably, infiltrated-PDTO exhibited increased levels of T effector, cell cycle, and myeloid inflammation, and reduced levels of angiogenesis and FAO/AMPK, relative to tumor tissue.

**Figure 4.**
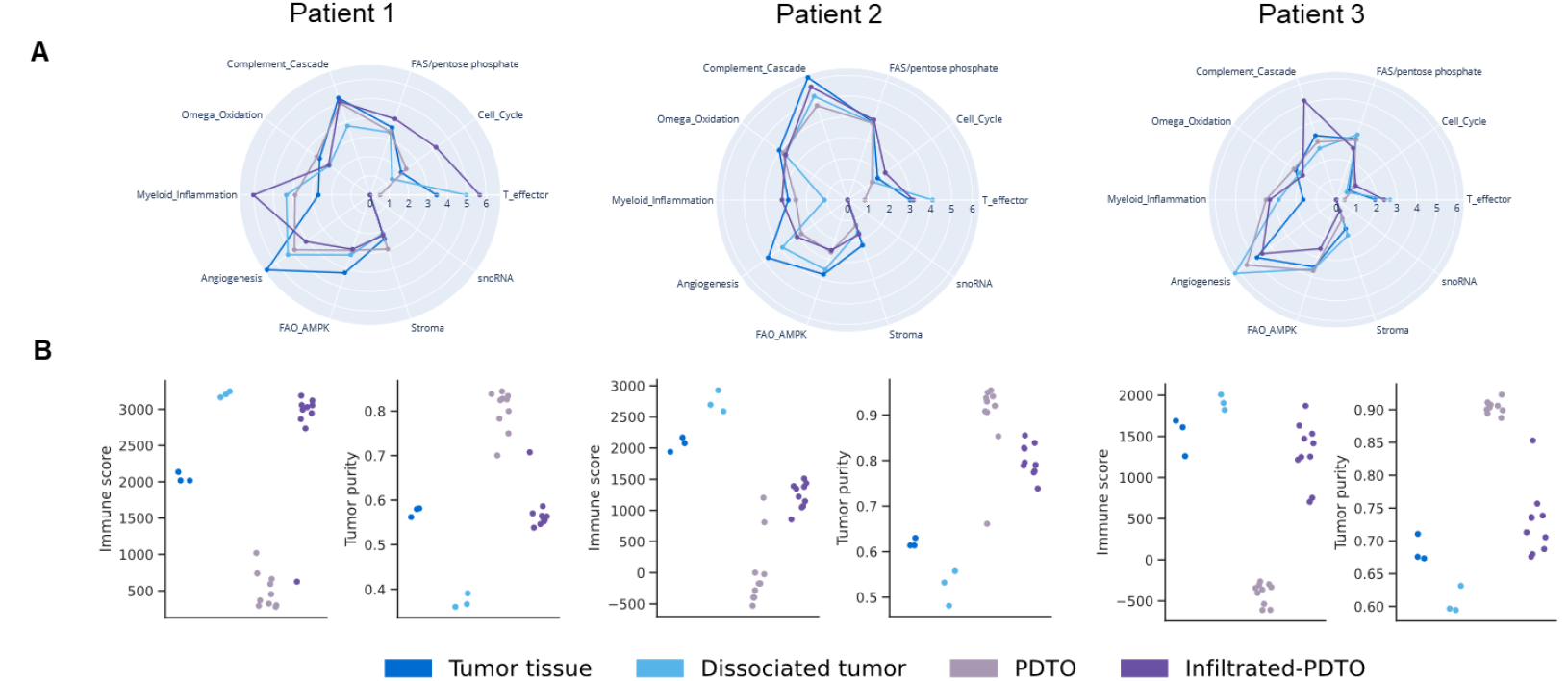
The signatures of cccRCC subtypes and EstimateScore of tumor tissue, dissociated tumor, PDTOs, and infiltrated-PDTOs from three patients. A. Ten simplified signatures of ccRCC subtypes. B. Immune score and tumor purity.

In Patient 2 and Patient 3, tumor tissues were characterized with the high level of complement cascade and angiogenesis, respectively. Consistently, dissociated tumor presented higher T effector than tumor tissue. While other signatures remain lower in dissociated tumor from Patient 2, myeloid inflammation, angiogenesis, and stromal cell signatures were higher in dissociated tumor from Patient 3. As the control, PDTO also showed the lowest level of T effector in both patients. Infiltrated-PDTOs maintained T effector signature levels comparable to tumor tissue and exhibited lower levels of angiogenesis and FAO/AMPK, despite differing subtypes. These may suggest the similar characteristic of TME and a restored metabolic reprogramming in infiltrated-PDTO to tumor tissue.

To assess tumor and immune cell composition, we employed the ESTIMATE algorithm to infer immune infiltration level and tumor purity^23^. Consistently, dissociated tumor had higher immune infiltration level (immune score) and lower proportion of tumor cells (tumor purity) than tumor tissue across three patients (Figure 4B; Table S8). While PDTO exhibited the lowest levels of immune cells due to the depletion of immune cells, infiltrated-PDTO displayed comparable levels of immune infiltration, and tumor percentage, compared to tumor tissue.

To accurately investigate which type of immune cells leads to the higher immune levels in infiltrated-PDTO, we performed single sample gene set enrichment analysis (ssGSEA) of 20 immune cells populations to quantify their infiltrating level in our samples (Figure 5; Table S9). Since PDTO was depleted of immune cells, the inferred infiltration levels of PDTO were taken as a control to adjust the immune infiltration level of other samples. In Patient 1, tumor tissue, dissociated tumor, and infiltrated-PDTO had similar infiltration levels of activated CD8 T cell, effector memory CD8 T cell, regulatory T cell, natural killer T cell, natural killer cell, myeloid-derived suppressor cell (MDSC), and eosinophil. Infiltrated-PDTO exhibited the decrease in activated B cell, activated dendritic cell, central memory CD8 T cell, and the total loss of effector memory CD4 T cell, gamma delta T cell, macrophage, mast cell, monocyte, neutrophil, immature dendritic cell (Figure 5). These suggest infiltrated-PDTO retained similar infiltration levels of key T cells, with the loss of myeloid cells.

**Figure 5.**
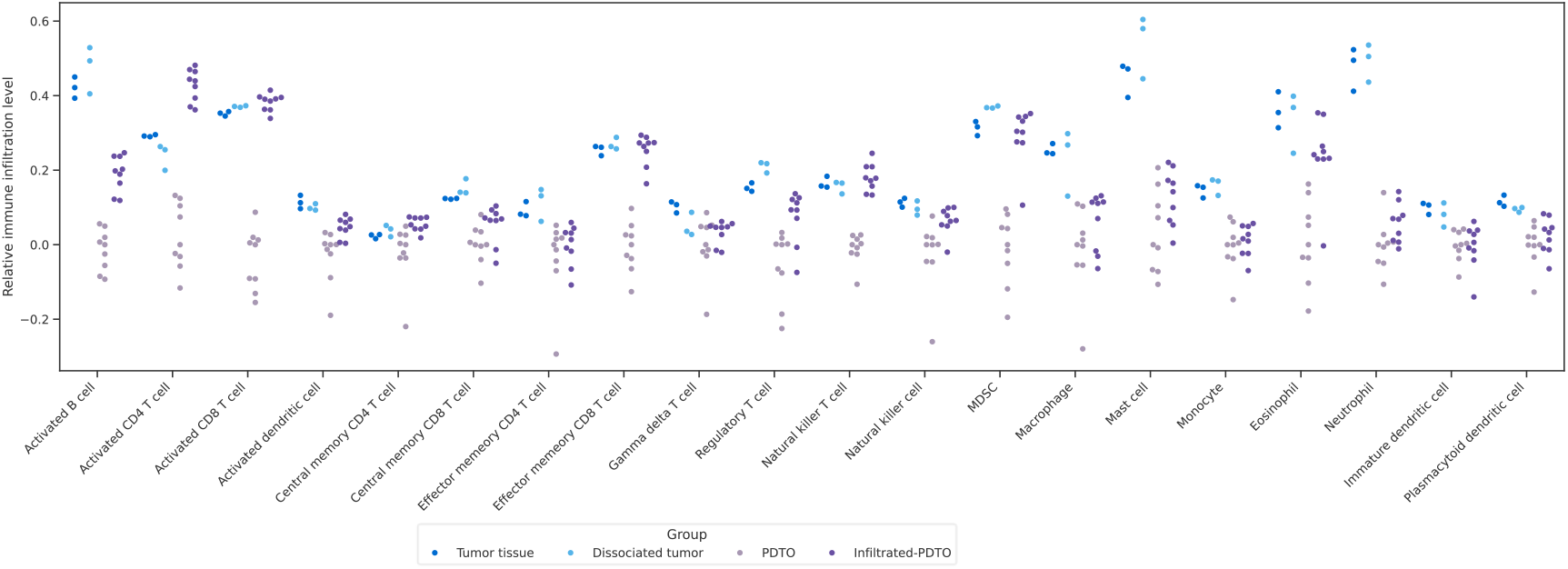
Immune infiltration level analysis of 20 immune cell types using single sample gene set enrichment analysis (ssGSEA) for Patient 1. GSEApy was employed to compute immune cell infiltration scores based on the transcriptomics of each sample. Since PDTOs were depleted of immune cells, the median of immune infiltration levels of PDTOs were taken as the control to adjust the estimated levels of immune cells in tumor tissue, dissociated tumor, and infiltrated-PDTOs.

In Patient 2 and 3, infiltrated-PDTO also exhibited comparable infiltration levels of activated CD8 T cell as tumor tissue showed (Figure S12). This was corroborated by similar immune scores between tumor tissue and infiltrated-PDTOs (Figure 4B). It was noted that infiltrated-PDTO samples from patient 3 exhibited significant variability in immune infiltration levels.

These findings underscore the critical role of immune cell re-infiltration in PDTOs, to accurately replicate the immune microenvironment of the original tumor. The observed variability of ccRCC subset signatures, tumor purity, and immune cell populations suggests these factors can influence the fidelity of infiltrated-PDTO, in modeling the original tumor’s characteristics. Therefore, meticulous control over tumor composition during organoid formation is essential to ensure that infiltrated-PDTOs serve as reliable models for studying tumor biology.

### Prediction of immunotherapy response (Figure 6)

To evaluate the potential of infiltrated-PDTO can be used as reliable models the therapeutic studies, we analyzed Tumor Immune Dysfunction and Exclusion (TIDE) scores, and expression of predictive genes across various samples. A lower TIDE score suggests a higher likelihood of positive response to immunotherapy, and samples with TIDE score below 0 were classified as responders. In Patient 1, both tumor tissue and infiltrated-PDTO samples exhibited comparable TIDE scores, with all samples (3/3 tumor tissue and 9/9 infiltrated-PDTO) classified as responders to immunotherapy (Figure 6; Table 14). In contrast, PDTO samples, which lack immune components, showed elevated TIDE scores and were uniformly labeled as non-responders. In Patient 2 and Patient 3, TIDE scores varied, resulting in 5 out of 10 (50%) and 2 out of 10 (20%) infiltrated-PDTO samples being classified as responders, respectively, compared to 1 out 3 (33.34%) in tumor tissue (Figure 6). These findings underscore the representativeness of infiltrated-PDTO in predicting immunotherapy response to the original tumor tissue.

**Figure 6.**
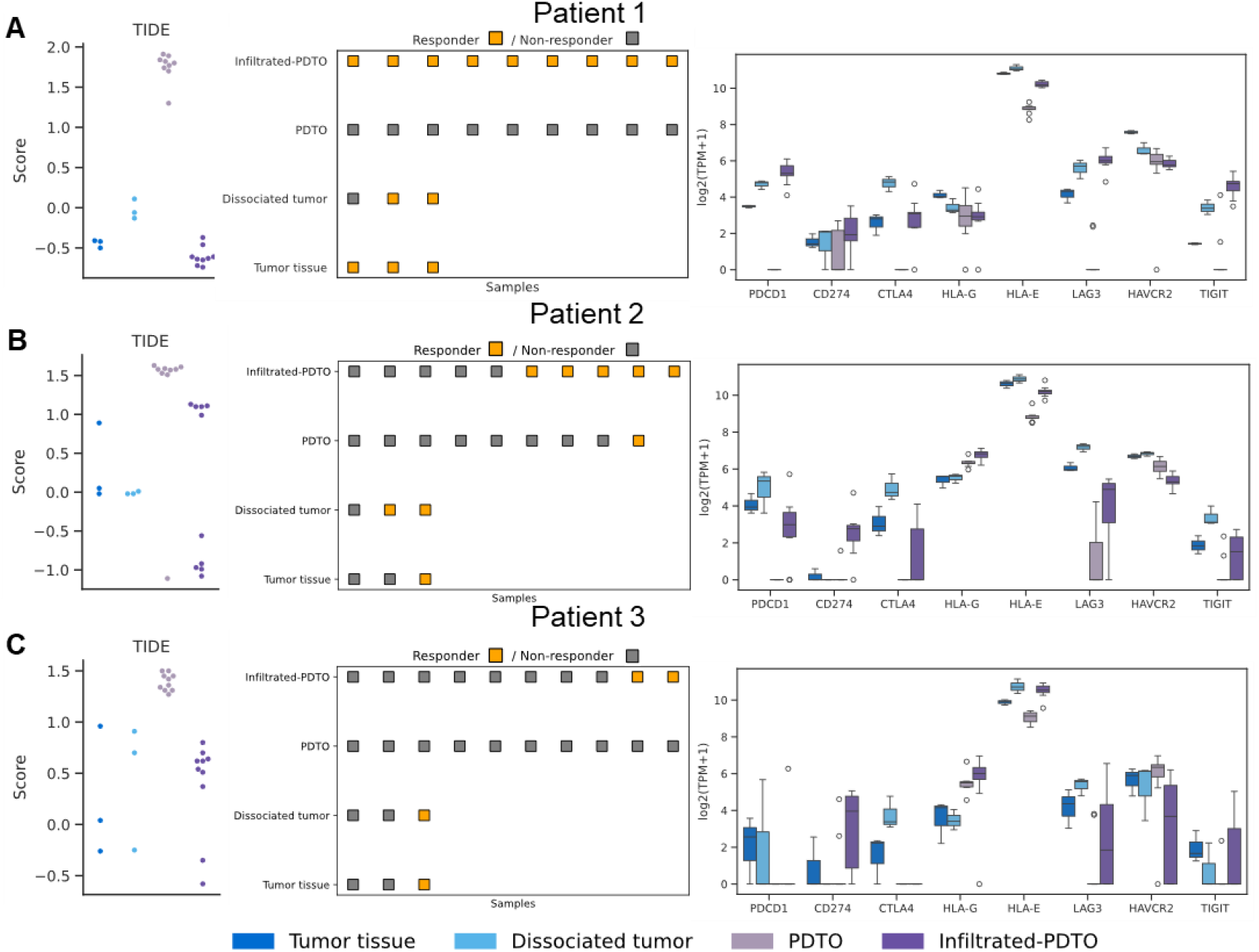
The TIDE scores, responder classification, and immune checkpoint genes expression for samples of tumor tissue, dissociated tumor, PDTOs, and infiltrated-PDTOs from three patients. TIDE scores were calculated through the web tool: http://tide.dfci.harvard.edu/. Sample were classified as responders if their TIDE score was below zero.

We further examined the expression of specific genes, including immune checkpoints (*PDCD1*: encoding PD-1, *CD274*: encoding PD-L1, *CTLA-4, HLA-G, LAG3, HAVACR2*: encoding Tim3, *TIGIT*), tumor-intrinsic mutators (*VHL, PBRM1, BAP1, SETD2, KDM5C*), and inflammatory genes (e.g., *CXCL9, CXCL10*) (Figure 6; Figure S13). In Patient 1, *PDCD1, CD274, CTLA4*, and *HLA-G* were expressed in tumor tissue, indicating tumor immune evasion, and potential resistance to immunotherapy. Infiltrated-PDTO demonstrated even higher expression levels of these genes, suggesting a restoration of immune suppression within the organoid model. *HLA-G* expression was slightly reduced, potentially reflecting differences in cell composition. *LAG3* and *TIGIT* expressed higher but *HAVCR2* expressed lower in infiltrated-PDTO compared to tumor tissue, possibly reflecting the early or intermediate exhaustion status of T cells in infiltrated-PDTO, as opposed to the terminal exhaustion status of T cells in tumor tissue. While tumor-intrinsic gene expression remained similar, suggesting retained mutation profile of infiltrated-PDTOs, inflammatory genes expressed higher in infiltrated-PDTOs, indicating the potential effect of IL-15 (Figure S13).

Compared with Patient 1, gene of PD-L1 was barely expressed in tumor tissue of Patient 2 and 3 (Figure 6), indicating less immune suppressive environment and poor response to immunotherapy. Infiltrated-PDTOs displayed lower expressions of *PDCD1* and *CTLA4*, but higher *CD274* expression. This could be IL-15, the cytokine, which activated immune cells and induce the expression of PD-L1 in infiltrated-PDTOs. For infiltrated-PDTOs, the lower expression of *HAVCR2* in Patient 2, and the slightly higher expression of *LAG3* and *TIGIT* in Patient 3, indicated the early or intermediate exhaustion status of T cells. Consistent with Patient 1, tumor-intrinsic genes remained unchanged between infiltrated-PDTOs and tumor tissue, and inflammatory genes expressed higher in infiltrated-PDTOs.

In summary, TIDE scores and immunotherapy markers suggested that infiltrated-PDTOs can serve as predictive models for immunotherapy response. While tumor-intrinsic gene expression remained stable, variations in immune checkpoint in infiltrated-PDTOs may influence their predictive accuracy for treatment response.

## Discussion & Conclusion

Patient derived organoids have become widely used as preclinical models in cancer research due to their ability to preserve tumor-specific characteristics. However, their representativeness is challenging to estimate, given experimental artifacts and the absence or alteration of the native tumor micro-immune environment. In this study, transcriptomic profiling across various stages of organoid formation was employed to dissect technical biases and patient-specific variability, offering critical guidance for the application of matched organoids in immunotherapeutic applications.

Tumor dissociation, as enzymatic digestion in our protocol, is an essential step to initiate organoids culture. Although it facilitates the isolation and manipulation of specific cell populations, it also disrupts essential features such as the extracellular matrix, spatial architecture, and native cellular interactions within the tumor^13^. Previous studies have demonstrated that dissociation process may trigger immediate early genes (IEGs), heat shock proteins, and stress response pathways^28^. Our findings align with this, revealing the up-regulation of ATP related genes (eg. *ATP5F1A*), ribosomal related genes (eg, *RPL10A*), heat shock related genes (eg. *HSPA1A*), and IEGs (eg. *JUNB*). Moreover, we observed the down-regulation of solute carrier family genes (eg. *SLC12A7*), ERBB3 signaling pathway, and hypoxia pathway, likely triggered by the loss of tumor structure. Notably, *SLC12A7* and *ERBB3* have been reported to be prognostic markers in ccRCC^31,32^, suggesting a disrupted role of dissociated tumor samples as potential models. Additionally, tumor dissociation may lead to the increase of the activity of immune cells, which could overestimate treatment response of tumors. These findings highlight biased transcriptomics following tumor dissociation and necessitate organoids formation in recapitulating key tumor structures and signaling pathways.

Post-dissociation, the generation, and analysis of both standard PDTOs and immune-infiltrated PDTOs revealed further insights into tumor reconstitution. In the absence of immune components (used as a control), PDTO restored hypoxia pathway and displayed down-regulation of immune cell markers, immune signatures, and inflammatory pathways In contrast, immune-infiltrated PDTOs displayed upregulation of proinflammatory genes such as *CXCL10* and *STAT1*, along with activation of the type II interferon signaling pathway ^33^ Simultaneously, there was downregulation of IEGS, and stress response genes in infiltrated-PDTOs relative to the dissociated tumor, indicating immune-mediated injury to organoids and reduced cellular stress. Importantly, robust activation of the JAK-STAT pathway was consistently observed in infiltrated-PDTOs across all three patients,, supporting the successful recruitment of immune cells into the organoid structure, and a partial restoration of tumor-immune crosstalk^34^.

The primary objective of this study was to evaluate the extent to which infiltrated-organoids can mimic the original tumor, and their potential for studying immunotherapy resistance. Our results indicate infiltrated-PDTO preserved several key immune signatures, including the T effector signature, immuneScore, and activated CD8+ T cell signatures, as well as comparable proportions of predicted responders inferred by TIDE score. These findings collectively suggest infiltrated-PDTOs are representative of the original tumor in the context of immunotherapeutic profiling. This representativeness may be attributed to the preservation of native immune components, reconstitution of the tumor microenvironment (TME), and structural resemblance to the parent tumor similarity^35,36^.

Nevertheless, several limitations must be acknowledged. First, the efficiency and fidelity of infiltrated organoids are influenced by inter-patient tumor heterogeneity. The number of DEGs varied largely across three patients, resulting in distinct TFs and pathway signature. For instance, PDTOs from Patient 1 displayed heightened myeloid inflammation, whereas those from Patient 3 showed enrichment in complement cascade pathways. T cell exhaustion markers also varied between patients, highlighting the intrinsic heterogeneity of ccRCC. Second, while IL-15 was employed to activate CD8^+^ T cells, promoting their proliferation, survival, and distribution^37^, its use carries limitations. The observed strong activation of the JAK-STAT pathway in infiltrated-PDTOs is consistent with IL-15 engagement. However, high concentrations of IL-15 may introduce cytokine-driven toxicity, checkpoint upregulation, and potentially distort the immune landscape compared to the native tumor environment. Third, critical components of the TME, particularly myeloid and endothelial cells, were absent in infiltrated-PDTOs. These cell populations are key regulators of immune cell behavior and T cell responsiveness. Their absence may limit the model’s accuracy in fully representing tumor-immune interactions. Future efforts to incorporate these missing cell types will be essential for enhancing the fidelity and translational relevance of infiltrated-PDTOs as immunotherapy testing platforms.

In conclusion, this study demonstrates that immune-infiltrated PDTOs can effectively preserve tumor-specific immune signatures and structural features, making them a promising model for investigating immune responses and therapy resistance in ccRCC. By systematically evaluating the effects of tumor dissociation, organoid formation, and immune cell reconstitution, we provide a framework for improving the biological fidelity of these models. Further refinement of this infiltrated-PDTO platform will advance personalized immunotherapy and translational cancer research.

## DATA AVAILABILITY

The data and codes were deposited in the GitHub repository (https://github.com/liangwei01/Patient-derived-tumor-organoids).

## AUTHOR CONTRIBUTION

LY, LL, JR, JL, and CB conceived the study. LL performed the experiments. LY developed and performed the analysis. LY, LL, JR, JL, and CB wrote the manuscript. LY, LL, JR, JL, and CB contributed to the revision of the manuscript.

## FUNDING

This study was supported by EDISCE grant from University of Grenoble Alpes. This work was supported by the KATY project, which has received funding from the European Union’s Horizon 2020 research and innovation program under grant agreement no. 101017453, by the CANVAS project, which has received funding from the Horizon Europe twinning program under grant agreement no. 101079510, and by the DIGPHAT project (Multi-scale and longitudinal data modeling in pharmacology: toward digital pharmacological twins), which has received funding from the French research initiative ‘‘France 2030’’ through the program PEPR Digital Health under ANR grant agreement no. 22-PESN-0017. This work was also supported by the IDEX outgoing international mobility grant of Universite’ Grenoble Alpes.

## ACKNOWLEDGMENTS

Part of the computations were carried out using the GRICAD infrastructure (https://gricad.univ-grenoble-alpes.fr), which is supported by Grenoble research communities.

## SUPPLEMENTARY MATERIAL

Supplementary tables and figures are available online.

## Notes

### Competing Interest Statement

The authors have declared no competing interest.

